# DrugOrchestra: Jointly predicting drug response, targets, and side effects via deep multi-task learning

**DOI:** 10.1101/2020.11.17.385757

**Authors:** Yuepeng Jiang, Stefano Rensi, Sheng Wang, Russ B. Altman

**Affiliations:** Department of Electrical and Computer Engineering, University of California, San Diego, USA; Department of Bioengineering, Stanford University, Stanford, CA, USA; Department of Genetics, Stanford University, Stanford, CA, USA

**Keywords:** Multi-task learning, drug target prediction, drug side effect prediction, drug response prediction

## Abstract

Massively accumulated pharmacogenomics, chemogenomics, and side effect datasets offer an unprecedented opportunity for drug response prediction, drug target identification and drug side effect prediction. Existing computational approaches limit their scope to only one of these three tasks, inevitably overlooking the rich connection among them. Here, we propose DrugOrchestra, a deep multi-task learning framework that jointly predicts drug response, targets and side effects. DrugOrchestra leverages pre-trained molecular structure-based drug representation to bridge these three tasks. Instead of directly fine-tuning on an individual task, DrugOrchestra uses deep multi-task learning to obtain a phenotype-based drug representation by simultaneously fine-tuning on drug response, target and side effect prediction. By coupling these three tasks together, DrugOrchestra is able to make predictions for unseen drugs by only knowing their molecular structures. We constructed a heterogeneous drug discovery dataset of over 21k drugs by integrating 8 datasets across three tasks. Our method obtained significant improvement in comparison to methods that were trained on a single task or a single dataset. We further revealed the transferability across 8 datasets and 3 tasks, providing novel insights for understanding drug mechanisms.

**Availability:** https://github.com/jiangdada1221/DrugOrchestra

## 1. Introduction

Large-scale pharmacogenomics studies[1–3], which investigate how genes affect an individual’s response to drugs, pave the path towards precision medicine[4,5]. Likewise, massively generated chemogenomics screens and patient side effect records have been used to identify drug-target interactions (DTIs) and adverse drug reactions (ADRs) in bulk. To date, more than 15 million DTIs and 0.6 million ADRs have been collected in public resources[6–10], presenting an unprecedented opportunity for drug discovery. In pursuit of this vision, computational approaches have been developed to predict drug response, targets, and side effects, which further prioritize experiments for biologists and support clinical decision making for doctors[10–17]. For example, Torng and Altman proposed a two-step graph convolutional neural network framework to learn high-quality protein pocket representations for drug-target interaction prediction[15]. Kim et al. formulated drug response prediction as a linear integer program and identified drug sensitivity subnetworks[16]. Luo et al. proposed a network-based approach that integrated a variety of biological features for drug target identification[17].

However, despite the encouraging performance of these existing methods on each individual task, none of them has considered jointly modeling all three tasks, inevitably overlooking connections among them. From a biological perspective, molecular and cellular targets serve as the basis of uncovering drug mechanisms[18,19], which are further used to explain systematic drug response variations[3,20]. Interactions between a drug and unintended receptors that are crucial in normal cellular functions can result in severe adverse drug reactions[21]. From a computational perspective, integrating and jointly analyzing datasets from all three tasks might boost the prediction performance by utilizing common patterns to alleviate overfitting. More importantly, molecular structure-based drug representation such as SMILES strings[22] have shown to be useful in all these tasks and can thus be used to transfer knowledge across them. Therefore, investigating these three tasks collectively might better decipher drug mechanisms and shed light on future drug development.

Inspired by the strong connection among these three tasks, we propose to jointly learn all three tasks using multi-task learning (MTL). The key idea of MTL is to solve multiple prediction tasks at the same time while automatically exploiting similarities and differences across tasks. Empowered by the advance of deep learning, MTL has achieved promising performance in computer vision[23], speech recognition[24] and natural language processing[25], in comparison to single task learning, which optimized each task separately. In biomedicine, MTL has also recently demonstrated exciting progress in drug discovery. Bharath and Steven et al. used MTL to model large-scale experiments on 40 million measurements of over 200 targets[26]. Qiao et al. applied an MTL model to a heterogeneous dataset integrated from 12 different individual cancer drug response datasets[27]. Kamran et al. utilized MTL to improve large-scale gene expression inference[28]. However, since these methods treated each drug as a single task when exploiting MTL, they required a substantial number of training samples for each drug and thus cannot be applied to new drugs.

In this paper, we proposed DrugOrchestra, a deep multi-task learning method that can simultaneously predict drug response, drug targets, and drug side effects. DrugOrchestra first used drug molecular structure-based drug representation pre-trained from millions of compounds[29] as shared features to bridge these three tasks. It then used a hard parameter sharing deep learning structure to jointly optimize all three tasks. Such structure enables DrugOrchestra to not only gain performance improvement on each single task but also make predictions for unseen drugs, which have never been seen in the training samples in any of these three tasks. This ability to predict for unseen drugs would play a key role in extending pharmacogenomics studies from thousands of tested compounds to millions of underexplored small molecules. We applied our method to 8 datasets across three tasks, including Repurposing Hub, Drugbank, STITCH, PDX, GDSC, CCLE, SIDER, and OFFSIDES, which in total cover more than 21k compounds. We observed significant improvement of our method on seven of these datasets. Our method further reveals the similarity and transferability across these datasets and across these tasks, providing novel insights for understanding drug mechanisms.

## 2. A heterogeneous drug discovery dataset

We first introduce the new heterogeneous drug discovery dataset we collected for drug response, targets and side effects prediction, including 21,032 compounds, 2,057 cell lines, 400 PDX models, 17,563 genes and 1,005 side effects.

### Drug target dataset

We obtained DTIs from three sources: Repurposing Hub[6], Drugbank[7] and STITCH[8]. Repurposing Hub contains 13,563 DTIs from 6,798 drugs and 2,183 gene targets. Drugbank contains 25,569 drug-target DTIs from 13580 drugs and 4673 gene targets. STITCH consists of 15,473,939 DTIs from 787,039 drugs and 19,195 gene targets. For each dataset, we excluded drugs whose SMILES strings were not in PubChem[30] to enable that each compound had a corresponding SMILES string representation. For the STITCH database, we excluded drug-target interactions that have a confidence score of less than 0.9. The number of drugs within each dataset and the overlap of drugs across datasets are shown in **Fig. 1a**. On average, there are 2.7, 3.6, and 3.1 gene targets associated with each drug and 5.7, 6.9, and 15.3 drugs associated with each target in Repurposing Hub, Drugbank, and STITCH, respectively.

**Fig. 1.**
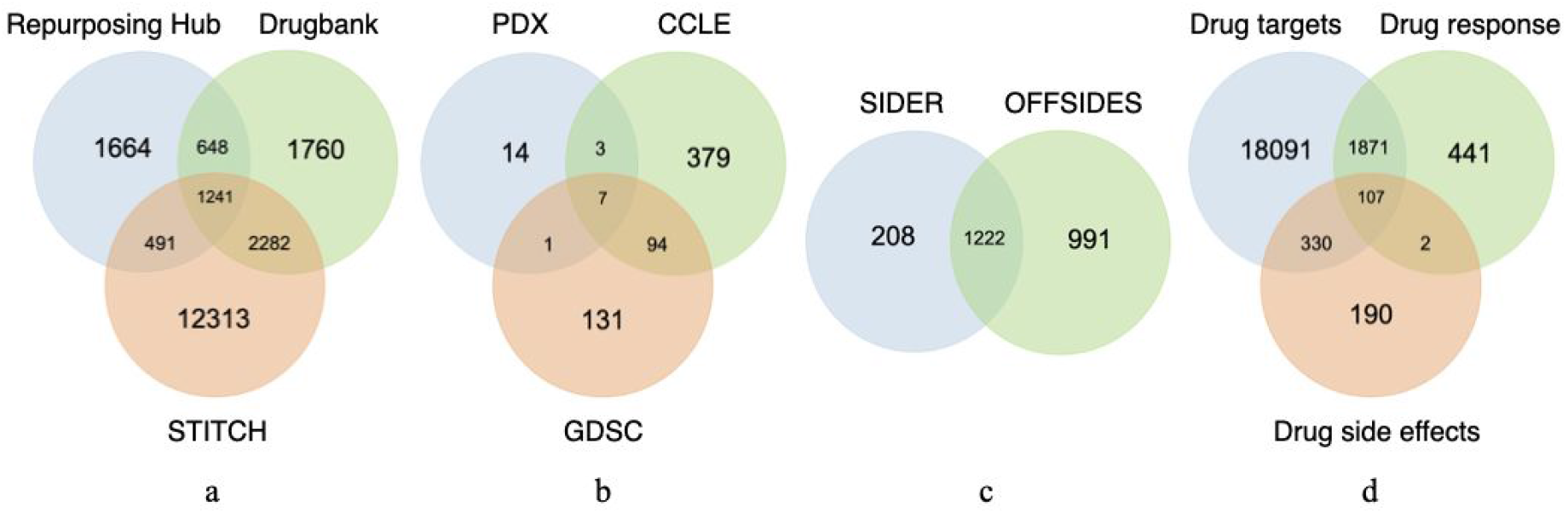
Overlap of drugs among different datasets and tasks. **a-c,** Venn diagrams showing the number of drugs within each dataset and the overlap of drugs among different datasets in drug targets (a), drug response (b) and drug side effects (c). **d,**Venn diagram showing the number of drugs with each task and the overlap of drugs among different tasks.

To construct the vector representations of drugs, we utilized the simplified molecular-input line-entry system (SMILES)[18], which uses a line notation to represent the structure of small molecules. In our paper, we used the pre-trained chemical molecular embedding model from Hu and Liu et al.[29], which takes SIMILES strings as inputs to obtain the drug representation. This pre-trained model was trained on millions of unlabeled chemical structure data, which enables the model to capture the domain-specific knowledge in the 2D molecular graph described by a SMILES string. We then obtained the low-dimensional drug features for each drug in the 8 datasets by inputting the corresponding SMILES string to the pre-trained model.

We obtained the target gene features from the pre-trained representations of genes in humans and several model organisms computed by the state-of-the-art network embedding method Mashup[31]. In brief, Mashup integrated multiple individual networks into a low-dimensional space according to the topological relationship between nodes. We mapped each gene in drug-target datasets to the gene vector in the pre-trained gene vector file provided by Mashup. We then excluded target genes that were not included in the gene list of Mashup. After the gene feature extraction, there are 2,183 target genes, 3,157 target genes, 17,288 target genes remained in Repurposing Hub, Drugbank, and STITCH respectively.

### Drug response dataset

We built a drug response collection by integrating a Patient-Derived Xenograft (PDX) dataset[32], the Genomics of Drug Sensitivity in Cancer (GDSC)[2] and the Cancer Therapeutics Response Portal (CCLE)[3]. The PDX dataset has 400 PDX model-derived tumor xenograft models with a diverse set of driver mutations. It consists of 37 drugs across 400 PDX model samples. The GDSC dataset has 255 drugs across 1018 cell lines. The CCLE dataset profiles 545 drugs across 1039 cell lines. We used gene expression profiles as the feature for drug response prediction. We used IC50 values as the indicator of the drug response data in GDSC and CCLE and Best Tumor Response value as the indicator of the drug response data in PDX. We also excluded drugs without the corresponding SMILES string in PubChem. For the response values, we took the z-score of the original response data for each dataset. After filtering, 1,634, 190,853, 322,045 drug response data points (i.e., cell line drug pairs) remained in PDX, GDSC, CCLE respectively. The number of drugs within each dataset and the overlap of drugs among three drug response datasets are shown in **Fig. 1b**.

We formed the cell line (PDX model) features based on the gene expression matrix which illustrates relationships between cell lines and genes. In the gene expression matrices of drug-response datasets, we only keep common genes that appear in all the three drug-response datasets. We then applied the PCA (Principal Component Analysis)[33] technique to the filtered gene expression matrices and obtained low-dimensional feature matrices. We took the z-score across rows to normalize the feature matrices. Each column in the feature matrices represents a feature vector for the corresponding cell line.

### Drug side effect dataset

We collected drug side effects from SIDER[9] and OFFSIDES[10]. The SIDER dataset contains 2,523,626 drug side-effect associations over 1,430 drugs and 5,868 side effects obtained by mining ADR events from drug label text. The OFFSIDES dataset consists of 0.32 million interactions between 2,730 drugs and 14,544 side effects collected from adverse event reporting systems. Likewise, we excluded drugs without the corresponding SMILES string in Pubchem. For OFFSIDES, we further excluded drug side-effect associations whose proportional reporting ratio errors are greater than 0.25. **Fig. 1c** shows the number of drugs within each dataset and the overlap of drugs between these two drug side effect datasets. On average, there are 73 and 108 diseases associated with each drug and 97 and 238 drugs associated with each disease in SIDER and OFFSIDES respectively.

We formulated the feature for a given disease by utilizing associations between diseases and genes pulled from DisGeNET[34]. The initial feature vector of each disease was an one hot vector, where 1 represented the existence of an association between the disease and the gene and 0 otherwise. Side effects that were not included in the DisGeNET as diseases were excluded. Following the same procedure of extracting cell line features, we used PCA to reduce the disease feature matrix (gene-by-disease) to a dimension of 300 and then applied z-score normalization to the low-dimensional vectors. 1,005 side effects remained after the processing and each column of the reduced feature matrix represented a feature vector for the corresponding disease. The overlap of drugs among three tasks and the statistics of each dataset are shown in **Fig. 1d** and **Table 1**.

**Table 1.**
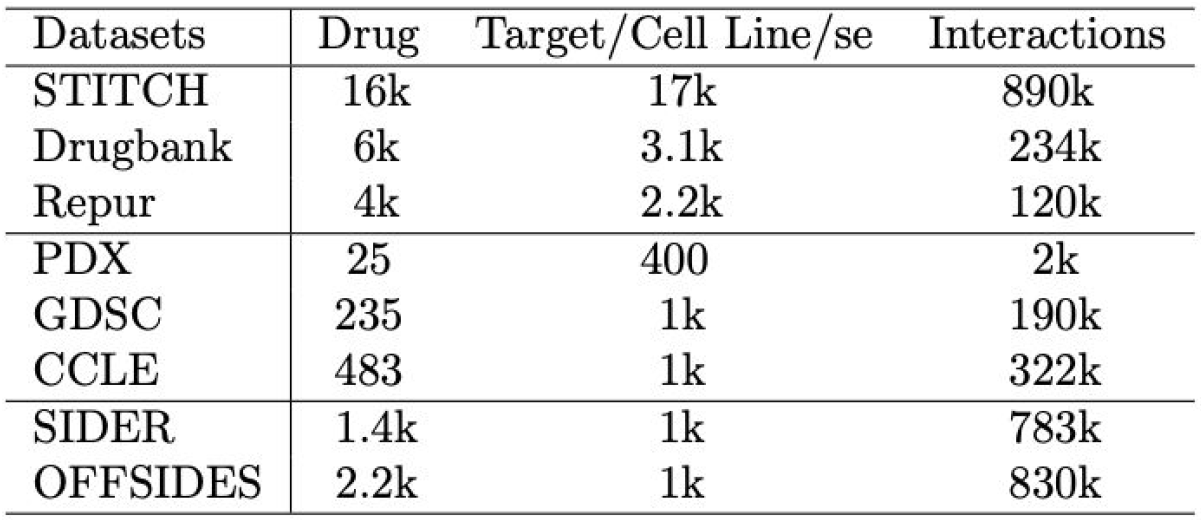
Statistics of the datasets included in our heterogeneous drug discovery dataset.

| Datasets | Drug | Target/Cell Line/se | Interactions |
| --- | --- | --- | --- |
| STITCH | 16k | 17k | 890k |
| Drugbank | 6k | 3.1k | 234k |
| Repur | 4k | 2.2k | 120k |
| PDX | 25 | 400 | 2k |
| GDSC | 235 | 1k | 190k |
| CCLE | 483 | 1k | 322k |
| SIDER | 1.4k | 1k | 783k |
| OFFSIDES | 2.2k | 1k | 830k |

## 3. Methods

### DrugOrchestra architecture

The structure of DrugOrchestra is shown in **Fig. 2**. Each task used a neural network as the base classifier. Each neural network consists of two components, shared layers across all tasks (three tasks in this paper) and task-specific layers. We used hard parameter sharing for parameters of the shared layers. Input features for shared layers are pre-trained molecular graph-based drug features. Input features for task-specific layers are different for each task. The output of the task-specific layers and the shared layers are concatenated and used to train the remaining layers of the task-specific neural network for each task.

**Fig. 2.**
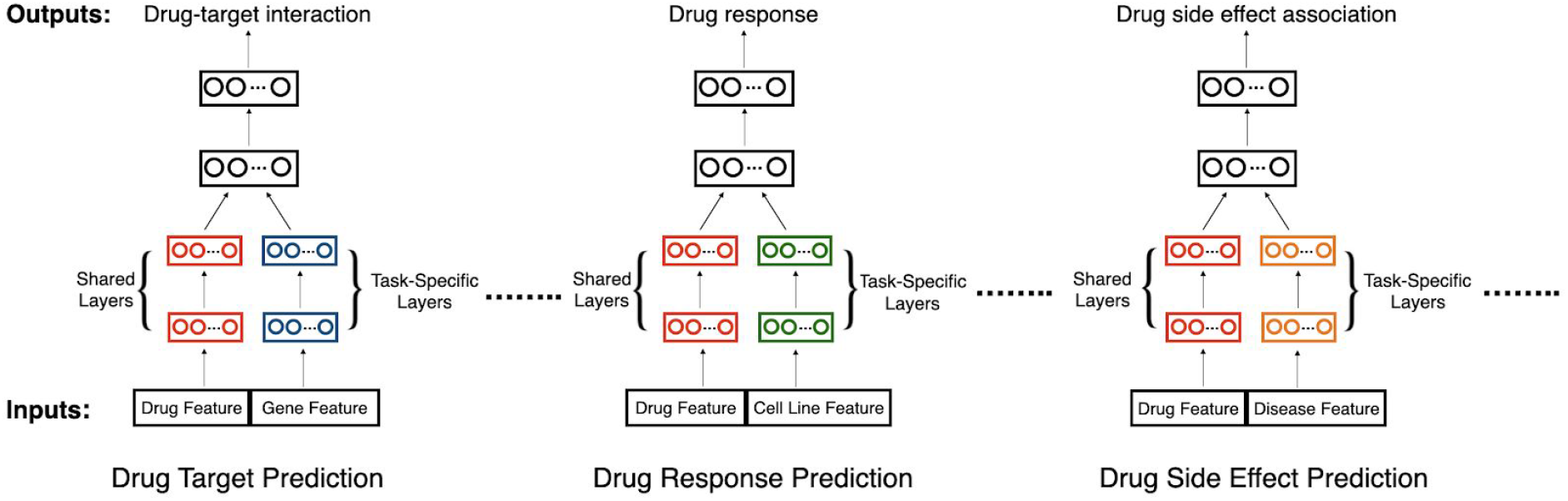
DrugOrchestra architecture. Drug features from each dataset are used by the same hidden layers using hard parameter sharing (red). Features other than drug features are used by task-specific layers (blue, green, orange). Outputs of these two layers are then concatenated as the input for the remaining task-specific neural networks (black).

In particular, let *d*_1_, *d*_2_…. *d_n_* denote input features for shared layers of *n* tasks. *f*_1_, *f*_2_….*f_n_* denote input features for task-specific layers of *n* tasks. For shared layers, the activations 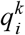of task *i* is calculated as:

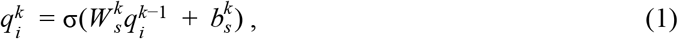

where the 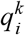 denotes the *k*-th shared hidden layer in the *i*-th task, σ is the activation function. 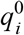 is the input drug feature *d*_*i*_. 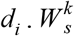 and 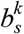 represent the weight and bias of the *k*-th hidden layer, which are shared among different tasks. Drug features from different datasets are processed by the same hidden layers to capture common patterns across tasks. We used the ReLU function as the activation function σ for hidden layers. We used the sigmoid function as the activation σ for the output layer of classification tasks (e.g., drug target prediction and drug side effect prediction) and the linear activation function for the activation σ of the output layer of the regression task (e.g., drug response prediction).

For the task-specific hidden layers that use task-specific features, the activations 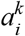 of each group is defined as:

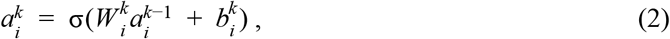

where 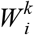 and 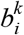 denote the weight and bias of the *k*-th task-specific hidden layer for the *i*-th task. 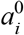 is the input task-specific feature *f_i_*. Different tasks can learn distinct feature representations which preserve the knowledge of each separate task. We used two hidden layers for both drug hidden layers and task-specific hidden layers. After we obtained the processed low-dimensional drug features and task-specific features, we concatenated them to get the new feature vector *c_i_* = [*q_i_*, *a_i_*] for task *i*. *c_i_* is then fed into a task-specific neural network with ReLU activation function. Let *o_i_* be the output value of the remaining task-specific neural network. *o_i_* = *sigmoid*(*c_i_*) for the classification task and *o_i_* = *c_i_* for the regression task.

Finally, we used the cross entropy loss for the classification task:

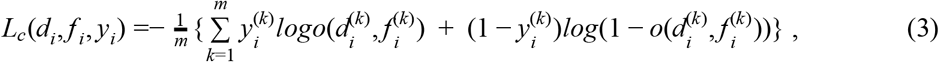

where m is the number of training sample in *i*-th task and 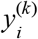 is the label of the *k*-th training sample in the *i*-th task. We used the mean squared error loss for the regression task:

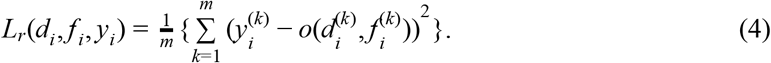

The overall objective for our multi-task learning model is to minimize the weighted combination of losses from each task, which is defined as:

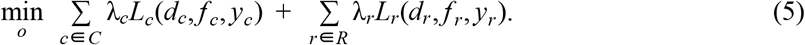

Here, *C* denotes a set of classification tasks, *R* denotes a set of regression tasks, λ*_i_* is the weight of task *i*. The DrugOrchestra framework is flexible to include more tasks that involve drug features. Through the multi-task learning framework, DrugOrchestra implicitly augments the training data and leverages other tasks to regularize a specific task, thus potentially alleviating overfitting.

### Dynamic Weight Adjustment

Training multiple tasks simultaneously could be challenging using multi-task learning since they need to be assigned proper importance weights. Extensively tuning hyperparameters for the weights of each task is time consuming. Therefore, we used a schema that can automatically adjust the weight of each task. In this paper, we compared two weight adjusting strategies introduced by Liu et al.[23] and Liu et al.[48] and observed that the dynamic weight adjustment strategy from Liu et al.[23] is empirically better. The dynamic weight adjustment (DWA) strategy first calculates the relative descending rate ω from two previous epochs:

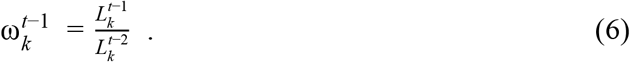

Here, *t* denotes the *t*-th epoch, *k* denotes the *k*-th task, and 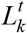 denotes the loss of the *k*-th task in the *t*-th epoch. Then the importance weight 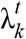 of task *k* in the *t*-th epoch is calculated as:

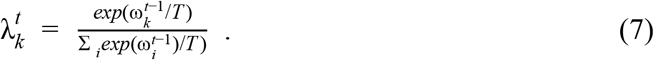

*T* controls the effect of task weighting. A smaller *T* makes the optimization process aggressively focused on important tasks determined by ω. This weight is used to weigh the loss function of each specific and DWA allocates higher weights to tasks with lower descending rates to boost the reduction of losses of those tasks. We set *T*=2 in our experiments as suggested by previous work[23].

## 4. Experimental setup

We compared our multi-task learning framework against three comparison approaches. 1) **Linear model**: We applied the Support Vector Machine[36] (SVM) with linear kernel to each single dataset. The learning rate schedule is set to be ‘optimal’ and the training stops until convergence. 2) **Ensemble model**: We applied the Random Forest[38] to each single dataset. The model was implemented using the sklearn library[37] with trees and depth set to 100 and 10 respectively. 3) **Single-task Learning**: We used a single-task deep neural network (STL-NN) model, which exploited the same architecture as DrugOrchestra but only trained on a single task. Although all these three comparison approaches are trained on a single task, they used all available training samples from the corresponding datasets. Moreover, all comparison approaches also used the same pre-trained molecular features, input features and data splits as DrugOrchestra. We used the same learning rate and number of epochs to train DrugOrchestra and STL-NN.

Drug target prediction and drug side effect prediction are classification tasks. Therefore, we used the area under curve for Receiver Operating Characteristic (AUROC) and the area under Precision Recall Curve (AUPRC) to evaluate the performance of them. Large AUROC and AUPRC indicate a better performance. Drug response prediction is a regression task. Consequently, we used the Spearman’s Correlation Coefficient (Spearman) and the Mean Squared Error (MSE) to evaluate the performance. A larger SCC value indicates a better performance whereas a smaller MSE indicates a better performance.

We defined the transferability between the source dataset (task) and the target dataset (task) using the performance gain:

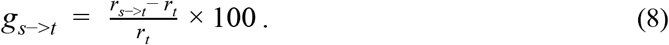

Here, *r_t_* is the performance of the target dataset *t*. *r*_*s*−>*t*_ represents the performance of training DrugOrchestra on the full source dataset *s* and the training set of dataset *t* jointly and then testing on the test set of dataset *t*. The Tanimoto score between two drugs are computed using the RDKit package[39]. A large Tanimoto score indicates that two drugs are similar in terms of the SMILES string.

We used 3-fold cross validation to train and evaluate our models. In particular, each time, we randomly selected 2/3 drugs for training and the remaining 1/3 drugs for testing. Since there is no overlap between test drugs and training drugs, this evaluation setting is able to examine whether our method can be used to predict for unseen drugs. We repeated this process for 10 times and used the average results as the final performance. The STL-NN model and DrugOrchestra were trained with ADAM optimizer[40] using a learning rate of 10^-3^, with a batch size of 256 for 20 epochs. The parameters of all layers were initialized by Xavier approach[49]. We used the binary cross-entropy (BCE) loss for drug-target prediction tasks and drug side effect prediction tasks, and the mean squared error (MSE) loss for drug response prediction tasks. The DrugOrchestra and the STL-NN model are implemented in Python using the Pytorch library[41].

## 5. Experimental results

### DrugOrchestra substantially improves performance on all three tasks

We first studied whether DrugOrchestra can improve prediction performance on these three tasks by comparing it with comparison approaches on 8 datasets of these 3 tasks. Results are summarized in **Table 2**. We found that 7 out of 8 datasets obtained significant improvement (t-test p-value<0.05). For example, in GDSC, DrugOrchestra achieved 0.375 SCC which is much higher than 0.315 SCC of STL-NN, 0.235 SCC of SVM and 0.149 SCC of Random Forest. Other six datasets also outperformed comparison methods significantly. The only exception is PDX where DrugOrchestra is better than RF but worse than SVM. We attributed the superior performance of linear SVM on PDX to the relatively smaller size of PDX, which could result in overfitting by using more expressive models. The overall improved performance of MTL demonstrates the effectiveness of using multi-task learning to transfer knowledge across these three tasks. Furthermore, we observed that DrugOrchestra had the most prominent improvement on drug response prediction, which might suggest a stronger connection between drug response and the other two tasks. Unlike the STL-NN model, SVM and Random Forest, DrugOrchestra models multiple datasets simultaneously so that it can take advantage of the shared latent features and obtain an enhanced drug representation.

**Table 2.** Comparison of DrugOrchestra with SVM, Random Forest and STL-NN on 8 datasets.

| Datasets | Metrics | SVM | Random Forest | STL-NN | DrugOrchestra |
| --- | --- | --- | --- | --- | --- |
| STITCH | AUROC | 0.668±0.048 | 0.66±0.032 | 0.827±0.031 | **0.833±0.047** |
| | AUPRC | 0.320±0.092 | 0.349±0.053 | 0.583±0.034 | **0.587±0.088** |
| Drugbank | AUROC | 0.602±0.042 | 0.616±0.037 | 0.740±0.034 | **0.752±0.012** |
| | AUPRC | 0.235±0.048 | 0.269±0.055 | 0.378±0.063 | **0.402±0.023** |
| Repur | AUROC | 0.530±0.010 | 0.585±0.049 | 0.691±0.014 | **0.703±0.031** |
| | AUPRC | 0.125±0.018 | 0.221±0.066 | 0.293±0.023 | **0.321±0.036** |
| PDX | SCC | **0.491±0.015** | 0.436±0.017 | 0.468±0.094 | 0.465±0.014 |
| | MSE | **0.824±0.034** | 0.864±0.034 | 0.856±0.028 | 0.840±0.031 |
| GDSC | SCC | 0.235±0.016 | 0.150±0.0140 | 0.315±0.020 | **0.375±0.024** |
| | MSE | 1.150±0.039 | 1.110±0.028 | 0.900±0.038 | **0.814±0.061** |
| CCLE | SCC | 0.206±0.016 | 0.181±0.012 | 0.394±0.022 | **0.410±0.010** |
| | MSE | 1.472±0.046 | 1.147±0.031 | 0.930±0.026 | **0.882±0.037** |
| SIDER | AUROC | 0.591±0.026 | 0.640±0.036 | 0.660±0.026 | **0.671±0.036** |
| | AUPRC | 0.231±0.022 | 0.295±0.035 | 0.312±0.016 | **0.321±0.034** |
| OFFSIDES | AUROC | 0.625±0.009 | 0.638±0.029 | 0.660±0.025 | **0.686±0.011** |
| | AUPRC | 0.399±0.005 | 0.435±0.013 | 0.432±0.022 | **0.457±0.037** |

### DrugOrchestra reveals transferability across datasets

The larger improvement on drug response prediction in comparison to the other two tasks motivates us to further investigate the transferability across datasets and tasks. To this end, we used performance gain to calculate the transferability across datasets and summarized the result in **Table 3**. In general, datasets belonging to the same task have better transferability compared to datasets from different tasks. For example, the average performance gain within drug response, targets and side effects prediction was 1.91%, 4.71% and 2.75% respectively, which is higher than the average performance gain 0.274%, 1.85% and 0.544% between datasets of different tasks. Datasets from the same task have closer distribution and the drug representation can thus be better shared across them. Moreover, we observed that the transferability can partially reflect the improvement of DrugOrchestra in **Table 2**. For instance, the GDSC dataset received the highest 18% improvement from CCLE dataset, and it also exhibited the highest improvement of 19% over STL. Interestingly, we also observed negative transferability in Table 3, which were mostly between datasets that were not from the same task. One exception was the transferability from PDX to CCLE, which again is consistent with the worse performance of DrugOrchestra on PDX in **Table 2**. We found that the remaining negative transferability was mostly between a drug side effect dataset and a drug target dataset, which might suggest the inherent difference between these two tasks.

**Table 3.** Transferability in terms of performance gain across datasets. Tasks in rows are source tasks and tasks in columns are target tasks. The source task is used to help the training of the target task. AUROC and SCC are used as evaluation metrics for the classification task (i.e., drug side effect prediction and drug target prediction) and the regression task (i.e., drug response prediction), respectively.

| Datasets | STITCH | Drugbank | Repur | PDX | GDSC | CTRP | SIDER | OFFSIDES |
| --- | --- | --- | --- | --- | --- | --- | --- | --- |
| STITCH | 0 | 2.414 | 2.356 | 1.399 | 3.700 | 1.149 | 0.953 | 0.543 |
| Drugbank | **1.090** | 0 | **2.642** | 0.9651 | 6.579 | 0.193 | 0.630 | 1.501 |
| Repur | 0.436 | **2.503** | 0 | 1.428 | 7.377 | 2.198 | 0.311 | 1.761 |
| PDX | -0.255 | 0.304 | 0.829 | 0 | 1.707 | -1.576 | -0.083 | -0.372 |
| GDSC | 0.004 | 0.504 | 0.679 | 2.221 | 0 | **7.261** | 0.111 | 0.382 |
| CTRP | 0.205 | 0.711 | 0.686 | 0.485 | **18.174** | 0 | 0.344 | 0.444 |
| SIDER | -0.628 | 1.128 | 0.482 | 1.670 | 3.318 | -2.928 | 0 | **3.733** |
| OFFSIDES | -0.427 | -0.027 | -0.084 | **2.251** | 2.342 | -3.970 | **1.767** | 0 |

### DrugOrchestra reveals transferability across tasks

Since the negative transferability is mostly observed between a drug side effect dataset and a drug target dataset, we then sought to examine the transferability at the task level. We performed the same analysis by aggregation datasets from the same task. **Table 4** shows the result of task level performance gain. We observed 4 out of 6 task pairs had positive performance gain by using DrugOrchestra. The improvement is most prominent when using drug targets to help the prediction of drug response. Drug target is known to be one of the most important features in a variety of drug response prediction approaches[42–44], partly due to its central role in understanding drug mechanisms. Interestingly, drug response is much less helpful for drug target prediction. The seemingly contradictory transferability might suggest the underlying causality relationship during apoptosis. Finally, we also observed substantial transferability between drug response and drug side effects, which are both drug phenotypes and have been shown to be associated[51]. The observed transferability at task levels raises our confidence that DrugOrchestra can be integrated with other datasets, which could further enhance the prediction performance on these three tasks.

**Table 4.** Transferability in terms of performance gain across tasks. Datasets belonging to the same task are combined as one dataset for that task. Tasks in rows are source tasks and tasks in columns are target tasks. The source task is used to help the training of the target task. AUROC and SCC are used as evaluation metrics for the classification task (i.e., drug side effect prediction and drug target prediction) and the regression task (i.e., drug response prediction), respectively.

| Tasks | Drug targets | Drug response | Drug side effects |
| --- | --- | --- | --- |
| Drug targets | 0 | 3.846 | -0.002 |
| Drug response | 0.006 | 0 | 0.276 |
| Drug side effects | -0.007 | 0.751 | 0 |

### DrugOrchestra enables prediction of unseen drugs

One key advantage of DrugOrchestra in comparison to other multi-task learning approaches[26–28] is its ability to make predictions for unseen drugs. Such improvement has been observed in **Table 2** where the test drug set has no overlap with the training drug set. However, compounds with similar molecular structures might still exhibit in the test and training set, which might cause potential information leakage and obscure the usage of our method in predicting unseen drugs. To investigate the applicability domain[50], we excluded a test set drug if it has a high Tanimoto similarity to any drug in the training set. By using different stringent Tanimoto similarity cutoffs, we still observed substantial improvement of DrugOrchestra in comparison to STL (**Table 5**). For example,when using a cutoff of 0.6, our method achieved 0.785 AUROC and 0.371 SCC in STITCH and GDSC respectively, which are much higher than 0.745 AUROC and 0.318 SCC of STL-NN. All the other 7 datasets including PDX obtained improved performance when using our method. Notably, the improvement of our method against STL was larger when using a more stringent cutoff, indicating the enhanced prediction performance of our method in making predictions for unseen drugs. By coupling these three tasks together and transferring knowledge across them, the new drug representation obtained by our method is more robust and can be generalized to unseen drugs.

**Table 5.** Performance of DrugOrchestra and STL-NN by only considering drugs above a specific Tanimoto threshold. AUROC is used for evaluating drug target datasets and drug side effect datasets, and SCC is used for evaluating drug response datasets.

| Methods(Threshold) | Datasets |  |  |  |  |  |  |  |
| --- | --- | --- | --- | --- | --- | --- | --- | --- |
|  | STITCH | Drugbank | Repur | PDX | GDSC | CCLE | SIDER | OFFSIDES |
| STL-NN(0.6) | 0.745±0.055 | 0.727±0.025 | 0.634±0.043 | 0.486±0.019 | 0.318±0.016 | 0.295±0.021 | 0.639±0.050 | 0.654±0.029 |
| DrugOrchestra(0.6) | **0.785±0.081** | **0.747±0.043** | **0.655±0.042** | **0.501±0.014** | **0.371±0.020** | **0.314±0.026** | **0.646±0.036** | **0.687±0.034** |
| STL-NN(0.7) | 0.775±0.038 | 0.732±0.034 | 0.651±0.043 | 0.468±0.020 | 0.312±0.014 | 0.338±0.014 | 0.643±0.049 | 0.654±0.024 |
| DrugOrchestra(0.7) | **0.806±0.053** | **0.746±0.052** | **0.668±0.021** | **0.477±0.016** | **0.380±0.016** | **0.358±0.018** | **0.648±0.030** | **0.689±0.023** |
| STL-NN(0.8) | 0.788±0.036 | 0.741±0.024 | 0.667±0.031 | 0.463±0.017 | 0.316±0.012 | 0.355±0.013 | 0.647±0.041 | 0.652±0.030 |
| DrugOrchestra(0.8) | **0.812±0.049** | **0.749±0.040** | **0.683±0.025** | **0.466±0.016** | **0.382±0.017** | **0.374±0.018** | **0.651±0.028** | **0.686±0.032** |
| STL-NN(0.9) | 0.806±0.047 | 0.751±0.020 | 0.679±0.031 | **0.467±0.088** | 0.322±0.078 | 0.373±0.011 | 0.651±0.039 | 0.654±0.031 |
| DrugOrchestra(0.9) | **0.827±0.027** | **0.756±0.038** | **0.692±0.030** | 0.462±0.015 | **0.388±0.018** | **0.394±0.016** | **0.658±0.027** | **0.687±0.031** |

## 6. Discussion

In this paper, we have presented DrugOrchestra, a novel deep multi-task learning model for jointly training drug response, targets, and side effects. DrugOrchestra exploits the task similarity among drug response, targets and side effects prediction. We collected a large-scale drug discovery dataset including 21,032 drugs, 17,563 genes, 2,057 cell lines and 1,005 side effects. Our method obtains significant improvement on 7 out of 8 datasets when evaluated on this dataset and thus enables more accurate predictions of response, side effects and targets for a given compound. We further showed the transferability across datasets and tasks, which were well aligned with our observation of the improvement and underlying drug mechanisms.

Our method is inspired by recent advances in the field of machine learning such as BERT[45] and GPT[46], which pre-trained on large-scale unlabeled data and then fine-tined on the specific task. Such two-stage approaches have been shown to be effective on a variety of tasks. A key conceptual advance of our method is to perform a multi-task learning step between the pre-training and fine-tuning step. In this paper, the multi-task learning is the joint training of drug response, side effects and targets. Based on the ablation study that compares to STL, we found that this multi-task learning step is crucial to the improvement, which suggests that knowledge of all three tasks can transfer to each other and enables us to get better molecular graph-based drug representation.

There are several directions for future research. From a biological perspective, we plan to integrate other tasks in the multi-task learning steps, such as drug perturbation prediction, drug solubility prediction and drug synergistic effect prediction. From a computational perspective, we would like to design interpretable methods to help us understand the multi-task learning model. In addition, we are also interested in experimentally validating the prioritized predictions by our model for drug repurposing and development.

## References

1. Barretina, Jordi, Giordano Caponigro, Nicolas Stransky, Kavitha Venkatesan, Adam A. Margolin, Sungjoon Kim, Christopher J. Wilson, et al. 2012. “The Cancer Cell Line Encyclopedia Enables Predictive Modelling of Anticancer Drug Sensitivity.” Nature 483 (7391): 603–7.

2. Yang, Wanjuan, Jorge Soares, Patricia Greninger, Elena J. Edelman, Howard Lightfoot, Simon Forbes, Nidhi Bindal, et al. 2013. “Genomics of Drug Sensitivity in Cancer (GDSC): A Resource for Therapeutic Biomarker Discovery in Cancer Cells.” Nucleic Acids Research 41 (Database issue): D955–61.

3. Rees, Matthew G., Brinton Seashore-Ludlow, Jaime H. Cheah, Drew J. Adams, Edmund V. Price, Shubhroz Gill, Sarah Javaid, et al. 2016. “Correlating Chemical Sensitivity and Basal Gene Expression Reveals Mechanism of Action.” Nature Chemical Biology 12 (2): 109–16.

4. Collins, Francis S., and Harold Varmus. 2015. “A New Initiative on Precision Medicine.” The New England Journal of Medicine 372 (9): 793–95.

5. Ashley, Euan A. 2016. “Towards Precision Medicine.” Nature Reviews. Genetics 17 (9): 507–22.

6. Corsello, Steven M., Joshua A. Bittker, Zihan Liu, Joshua Gould, Patrick McCarren, Jodi E. Hirschman, Stephen E. Johnston, et al. 2017. “The Drug Repurposing Hub: A next-Generation Drug Library and Information Resource.” Nature Medicine 23 (4): 405–8.

7. Wishart, David S., Yannick D. Feunang, An C. Guo, Elvis J. Lo, Ana Marcu, Jason R. Grant, Tanvir Sajed, et al. 2018. “DrugBank 5.0: A Major Update to the DrugBank Database for 2018.” Nucleic Acids Research 46 (D1): D1074–82.

8. Szklarczyk, Damian, Alberto Santos, Christian von Mering, Lars Juhl Jensen, Peer Bork, and Michael Kuhn. 2016. “STITCH 5: Augmenting Protein-Chemical Interaction Networks with Tissue and Affinity Data.” Nucleic Acids Research 44 (D1): D380–84.

9. Kuhn, Michael, Ivica Letunic, Lars Juhl Jensen, and Peer Bork. 2016. “The SIDER Database of Drugs and Side Effects.” Nucleic Acids Research 44 (D1): D1075–79.

10. Tatonetti, Nicholas P., Patrick P. Ye, Roxana Daneshjou, and Russ B. Altman. 2012. “Data-Driven Prediction of Drug Effects and Interactions.” Science Translational Medicine 4 (125): 125ra31.

11. Costello, James C., Laura M. Heiser, Elisabeth Georgii, Mehmet Gönen, Michael P. Menden, Nicholas J. Wang, Mukesh Bansal, et al. 2014. “A Community Effort to Assess and Improve Drug Sensitivity Prediction Algorithms.” Nature Biotechnology 32 (12): 1202–12.

12. Wang, Sheng, and Jian Peng. 2017. “Network-Assisted Target Identification for Haploinsufficiency and Homozygous Profiling Screens.” PLoS Computational Biology 13 (6): e1005553.

13. Tatonetti, Nicholas P., Tianyun Liu, and Russ B. Altman. 2009. “Predicting Drug Side-Effects by Chemical Systems Biology.” Genome Biology 10 (9): 238.

14. White, Ryen W., Sheng Wang, Apurv Pant, Rave Harpaz, Pushpraj Shukla, Walter Sun, William DuMouchel, and Eric Horvitz. 2016. “Early Identification of Adverse Drug Reactions from Search Log Data.” Journal of Biomedical Informatics 59 (February): 42–48.

15. Torng, Wen, and Russ B. Altman. 2019. “Graph Convolutional Neural Networks for Predicting Drug-Target Interactions.” Journal of Chemical Information and Modeling 59 (10): 4131–49.

16. Kim, Yoo-Ah, Rebecca Sarto Basso, Damian Wojtowicz, Dorit S. Hochbaum, Fabio Vandin, and Teresa M. Prztycka. 2019. “Identifying Drug Sensitivity Subnetworks with NETPHIX.” https://doi.org/10.1101/543876.

17. Luo, Yunan, Xinbin Zhao, Jingtian Zhou, Jinglin Yang, Yanqing Zhang, Wenhua Kuang, Jian Peng, Ligong Chen, and Jianyang Zeng. 2017. “A Network Integration Approach for Drug-Target Interaction Prediction and Computational Drug Repositioning from Heterogeneous Information.” Nature Communications 8 (1): 573.

18. Santos, Rita, Oleg Ursu, Anna Gaulton, A. Patrícia Bento, Ramesh S. Donadi, Cristian G. Bologa, Anneli Karlsson, et al. 2017. “A Comprehensive Map of Molecular Drug Targets.” Nature Reviews. Drug Discovery 16 (1): 19–34.

19. Schenone, Monica, Vlado Dančík, Bridget K. Wagner, and Paul A. Clemons. 2013. “Target Identification and Mechanism of Action in Chemical Biology and Drug Discovery.” Nature Chemical Biology 9 (4): 232–40.

20. Seashore-Ludlow, Brinton, Matthew G. Rees, Jaime H. Cheah, Murat Cokol, Edmund V. Price, Matthew E. Coletti, Victor Jones, et al. 2015. “Harnessing Connectivity in a Large-Scale Small-Molecule Sensitivity Dataset.” Cancer Discovery 5 (11): 1210–23.

21. Coleman, Jamie J., and Sarah K. Pontefract. 2016. “Adverse Drug Reactions.” Clinical Medicine 16 (5): 481–85.

22. Weininger, David. 1988. “SMILES, a Chemical Language and Information System. 1. Introduction to Methodology and Encoding Rules.” Journal of Chemical Information and Modeling 28 (1): 31–36.

23. Liu, S., E. Johns, and A. J. Davison. 2019. “End-To-End Multi-Task Learning With Attention.” In 2019 IEEE/CVF Conference on Computer Vision and Pattern Recognition (CVPR), 1871–80.

24. Deng, L., G. Hinton, and B. Kingsbury. 2013. “New Types of Deep Neural Network Learning for Speech Recognition and Related Applications: An Overview.” In 2013 IEEE International Conference on Acoustics, Speech and Signal Processing, 8599–8603.

25. Collobert, Ronan, and Jason Weston. 2008. “A Unified Architecture for Natural Language Processing: Deep Neural Networks with Multitask Learning.” In Proceedings of the 25th International Conference on Machine Learning, 160–67. ICML’08. New York, NY, USA: Association for Computing Machinery.

26. Ramsundar, Bharath, Steven Kearnes, Patrick Riley, Dale Webster, David Konerding, and Vijay Pande. 2015. “Massively Multitask Networks for Drug Discovery.” arXiv [stat.ML]. arXiv. http://arxiv.org/abs/1502.02072.

27. Liao, Qing, Ye Ding, Zoe L. Jiang, Xuan Wang, Chunkai Zhang, and Qian Zhang. 2019. “Multi-Task Deep Convolutional Neural Network for Cancer Diagnosis.” Neurocomputing 348 (July): 66–73.

28. Dizaji, Kamran Ghasedi, Wei Chen, and Heng Huang. 2020. “Deep Large-Scale Multi-Task Learning Network for Gene Expression Inference.” In Research in Computational Molecular Biology, 19–36. Springer International Publishing.

29. Hu*, Weihua, Bowen Liu*, Joseph Gomes, Marinka Zitnik, Percy Liang, Vijay Pande, and Jure Leskovec. 2020. “Strategies for Pre-Training Graph Neural Networks.” In International Conference on Learning Representations. https://openreview.net/forum?id=HJlWWJSFDH.

30. Kim, Sunghwan, Jie Chen, Tiejun Cheng, Asta Gindulyte, Jia He, Siqian He, Qingliang Li, et al. 2019. “PubChem 2019 Update: Improved Access to Chemical Data.” Nucleic Acids Research 47 (D1): D1102–9.

31. Cho, Hyunghoon, Bonnie Berger, and Jian Peng. 2016. “Compact Integration of Multi-Network Topology for Functional Analysis of Genes.” Cell Systems 3 (6): 540–48.e5.

32. Gao, Hui, Joshua M. Korn, Stéphane Ferretti, John E. Monahan, Youzhen Wang, Mallika Singh, Chao Zhang, et al. 2015. “High-Throughput Screening Using PDX model-Derived Tumor Xenografts to Predict Clinical Trial Drug Response.” Nature Medicine 21 (11): 1318–25.

33. Jolliffe, I. T. 1986. “Principal Component Analysis and Factor Analysis.” In Principal Component Analysis, edited by I. T. Jolliffe, 115–28. New York, NY: Springer New York.

34. Piñero, Janet, Juan Manuel Ramírez-Anguita, Josep Saüch-Pitarch, Francesco Ronzano, Emilio Centeno, Ferran Sanz, and Laura I. Furlong. 2020. “The DisGeNET Knowledge Platform for Disease Genomics: 2019 Update.” Nucleic Acids Research 48 (D1): D845–55.

35. Agarap, Abien Fred. 2018. “Deep Learning Using Rectified Linear Units (ReLU).” arXiv [cs.NE]. arXiv. http://arxiv.org/abs/1803.08375.

36. Cortes, Corinna, and Vladimir Vapnik. 1995. “Support-Vector Networks.” Machine Learning 20 (3): 273–97.

37. Pedregosa, Fabian, Gaël Varoquaux, Alexandre Gramfort, Vincent Michel, Bertrand Thirion, Olivier Grisel, Mathieu Blondel, et al. 2011. “Scikit-Learn: Machine Learning in Python.” Journal of Machine Learning Research: JMLR 12 (85): 2825–30.

38. Breiman, Leo. 2001. “Random Forests.” Machine Learning 45 (1): 5–32.

39. Landrum, Greg, and Others. 2006. “RDKit: Open-Source Cheminformatics.”

40. Kingma, Diederik P., and Jimmy Ba. 2014. “Adam: A Method for Stochastic Optimization.” arXiv [cs.LG]. arXiv. http://arxiv.org/abs/1412.6980.

41. Paszke, Adam, Sam Gross, Francisco Massa, Adam Lerer, James Bradbury, Gregory Chanan, Trevor Killeen, et al. 2019. “PyTorch: An Imperative Style, High-Performance Deep Learning Library.” In Advances in Neural Information Processing Systems 32, edited by H. Wallach, H. Larochelle, A. Beygelzimer, F. d\textquotesingle Alché-Buc, E. Fox, and R. Garnett, 8024–35. Curran Associates, Inc.

42. Zhang, Fei, Minghui Wang, Jianing Xi, Jianghong Yang, and Ao Li. 2018. “A Novel Heterogeneous Network-Based Method for Drug Response Prediction in Cancer Cell Lines.” Scientific Reports 8 (1): 3355.

43. Adam, George, Ladislav Rampášek, Zhaleh Safikhani, Petr Smirnov, Benjamin Haibe-Kains, and Anna Goldenberg. 2020. “Machine Learning Approaches to Drug Response Prediction: Challenges and Recent Progress.” NPJ Precision Oncology 4 (June): 19.

44. Menden, Michael P., Dennis Wang, Mike J. Mason, Bence Szalai, Krishna C. Bulusu, Yuanfang Guan, Thomas Yu, et al. 2019. “Community Assessment to Advance Computational Prediction of Cancer Drug Combinations in a Pharmacogenomic Screen.” Nature Communications 10 (1): 2674.

45. Devlin, Jacob, Ming-Wei Chang, Kenton Lee, and Kristina Toutanova. 2019. “BERT: Pre-Training of Deep Bidirectional Transformers for Language Understanding.” In Proceedings of the 2019 Conference of the North American Chapter of the Association for Computational Linguistics: Human Language Technologies, Volume 1 (Long and Short Papers), 4171–86. Minneapolis, Minnesota: Association for Computational Linguistics.

46. Chen, Mark, Alec Radford, Rewon Child, Jeff Wu, Heewoo Jun, Prafulla Dhariwal, David Luan, and Ilya Sutskever. 2020. “Generative Pretraining from Pixels.” In Proceedings of the 37th International Conference on Machine Learning.

47. Rumelhart, David E., Geoffrey E. Hinton, and Ronald J. Williams. 1986. “Learning Representations by Back-Propagating Errors.” Nature 323 (6088): 533–36.

48. S. Liu, Y. Liang, and A. Gitter, “Loss-balanced task weighting to reduce negative transfer in multi-task learning,” in Proc. of the 33rd AAAI Conf. on Artificial Intelligence, vol. 33, Honolulu, Hawaii, Jan. 2019, pp. 9977–9978.

49. Glorot, Xavier, and Yoshua Bengio. 2010. “Understanding the Difficulty of Training Deep Feedforward Neural Networks.” In Proceedings of the Thirteenth International Conference on Artificial Intelligence and Statistics, 249–56.

50. Ruiz, Irene Luque, and Miguel Ángel Gómez-Nieto. 2018. “Study of the Applicability Domain of the QSAR Classification Models by Means of the Rivality and Modelability Indexes.” Molecules 23 (11). https://doi.org/10.3390/molecules23112756.

51. Yang, Lun, and Pankaj Agarwal. 2011. “Systematic Drug Repositioning Based on Clinical Side-Effects.” PloS One 6 (12): e28025.

